# Long-term culture of fetal monocyte precursors *in vitro* allowing the generation of *bona fide* alveolar macrophages *in vivo*

**DOI:** 10.1101/2021.06.04.447115

**Authors:** Fengqi Li, Katarzyna Maria Okreglicka, Lea Maria Pohlmeier, Christoph Schneider, Manfred Kopf

## Abstract

Tissue-resident macrophage-based immune therapies have been proposed for various diseases. However, generation of sufficient numbers that possess tissue-specific functions remains a major handicap. Here, we show that fetal liver monocytes (FLiMo) cultured with GM-CSF (also known as CSF2) rapidly differentiate into a long-lived, homogeneous alveolar macrophage (AM)-like population *in vitro*. CSF2-cultured FLiMo remain the capacity to develop into *bona fide* AM upon transfer into *Csf2ra*^*-/-*^ neonates and prevent development of alveolar proteinosis and efferocytosis of apoptotic cells for at least 1 year *in vivo*. Compared to transplantation of AM-like cells derived from bone marrow macrophages (BMM), CSF2-cFliMo more efficiently engraft empty AM niches in the lung and protect mice from respiratory viral infection. Harnessing the potential of this approach for gene therapy, we restored a disrupted Csf2ra gene in FLiMo and their capacity to develop into AM *in vivo*. Together, we provide a novel platform for generation of immature AM-like precursors amenable for genetic manipulation, which will be useful to study to dissect AM development and function and pulmonary transplantation therapy.

## Introduction

Tissue-resident macrophages (MFTR) are heterogeneous cell populations, present in almost all tissues and play multiple tissue-specific functions in homeostasis and diseases (Davies et al, 2013; Hoeffel & Ginhoux, 2015). MΦ-based therapies have been proposed as potential strategies in various diseases (Duan & Luo, 2021; Mass & Lachmann, 2021; Moroni et al, 2019; Peng et al, 2020).

Lung resident AM play important roles in host defense to pulmonary infections and non-inflammatory clearance of inhaled particles (Hussell & Bell, 2014). Moreover, AM are required for catabolism of surfactant and removal of apoptotic cells and cellular debris that otherwise accumulate in the alveoli resulting in impaired air exchange (Kopf et al, 2015). AM are derived from late EMP and fetal liver monocytes, which colonize the lung during late embryogenesis, and are maintained life-long by self-renewal after birth (Hashimoto et al, 2013; Yona et al, 2013). Development and expansion are driven perinatally by GM-CSF-induced PPARγ (Guilliams et al, 2013; Schneider et al, 2014). Accordingly, absence of PPARγ, GM-CSF or one of the GM-CSFR subunits, Csf2ra and Csf2rb, in mice results in abrogated AM development and pulmonary alveolar proteinosis (PAP) (Guilliams et al, 2013; Nishinakamura et al, 1995; Robb et al, 1995; Schneider et al, 2017; Schneider et al, 2014). Moreover, human hereditary pulmonary alveolar proteinosis (herPAP) mutations, which is associated with long-term respiratory insufficiency and high susceptibility to microbial infection, is caused by mutations in *CSF2RA* and *CSF2RB* genes (Suzuki et al, 2011; Suzuki et al, 2008; Trapnell et al, 2003).

It has been shown that pulmonary transplantation of AM-like cells from BMM or induced pluripotent stem cells (iPSC) from yolk sac primitive macrophages or human blood CD34^+^ cells to *Csf2rb*^*-/-*^ mice can prevent development of PAP (Happle et al, 2018; Happle et al, 2014; Lachmann et al, 2015; Mucci et al, 2018; Suzuki et al, 2014; Takata et al, 2017). However, whether these transplanted MΦ resemble *bona fide* self-renewing AM with functional capacities beyond surfactant clearance remains unclear. Here, we have established a biologically relevant model to study AM development and function by gene editing *in vitro* and assessment of functional consequences *in vivo*. By culturing purified fetal liver monocytes with GM-CSF, we were able to generate high numbers of AM-like cells that can be kept proliferating like stem cells for several months *in vitro* and possess the ability to terminally differentiate and restore AM development and function upon pulmonary transplantation to *Csf2ra*^*-/-*^ neonates. CSF2-cFLiMo-derived AM resemble *bona fide* AM in gene expression profile, surface markers, and functional capacities including potent self-renewal capacity, clearance of surfactant, and efferocytosis of apoptotic cells during influenza virus infection. In contrast to BMM, CSF2-cFliMo more efficiently engraft empty AM niches and protect mice from influenza infection. Finally, CSF2RA re-expression in *Csf2ra*^*-/-*^ FLiMo by retroviral gene transfer restored their potential to develop into *bona fide* AM. Altogether, this platform provides a potent platform to study genes involved in AM development and function, as well as for therapeutic approaches.

## Results

### *In vitro* GM-CSF-differentiated fetal liver monocytes give rise to self-renewing cells with AM-like phenotype

The fetal liver contains several myeloid precursors including primitive macrophages (pMΦ) and EMP-derived monocytes (Gomez Perdiguero et al, 2015; Hoeffel et al, 2015). Recently, we and others have shown that the latter are the most potent precursors of AM (Li et al, 2020; van de Laar et al, 2016). To identify and better characterize the AM precursor, we sorted populations of viable F4/80^lo^CD11b^int^Ly6C^+^ monocytes and viable F4/80^hi^CD11b^lo^ pMF from the fetal liver of E14.5 C57BL/6 embryos (Fig. E1A) and cultured them with GM-CSF *in vitro* (Fig. 1A). Monocytes grew exponentially, whereas pMΦ disappeared within 2 weeks (Fig. E1B). At day 3 of the monocyte culture, a new population with macrophage characteristics (F4/80^+^Ly6C^-^ cells) emerged, which quickly predominated and appeared to be homogenous after 9 days in the culture (Fig. 1B). The monocyte-to-macrophage differentiation was paralleled by an increase in cell granularity (SSC) and expression of the surface markers CD11c and Siglec-F (Fig. E1C). Similar phenotypic changes also occur during fetal monocyte to AM differentiation in the lung, as described previously (Guilliams et al, 2013; Schneider et al, 2014), although high expression of CD11b indicated that cultured cells did not fully complete AM differentiation (Fig. E1D) (Schneider et al, 2014). Notably, the AM-like phenotype was maintained for at least 4 months of continuous culture (Fig. E1D). From here on, we refer to F4/80^+^CD11c^+^CD11b^+^Siglec-F^+^ cells as CSF2-cultured fetal liver monocytes (CSF2-cFliMo). Similar results were obtained when CSF2-cFLiMo were generated from monocytes isolated from E16.5, E18.5, and E20.5 fetal livers (Fig. E1E). When differentiated CSF2-cFliMo cells were deprived of GM-CSF, their numbers rapidly declined (Fig. E1F, G), although the remaining cells maintained the expression of AM surface markers including CD11c and Siglec-F (Fig. E1H).

**Figure 1.**
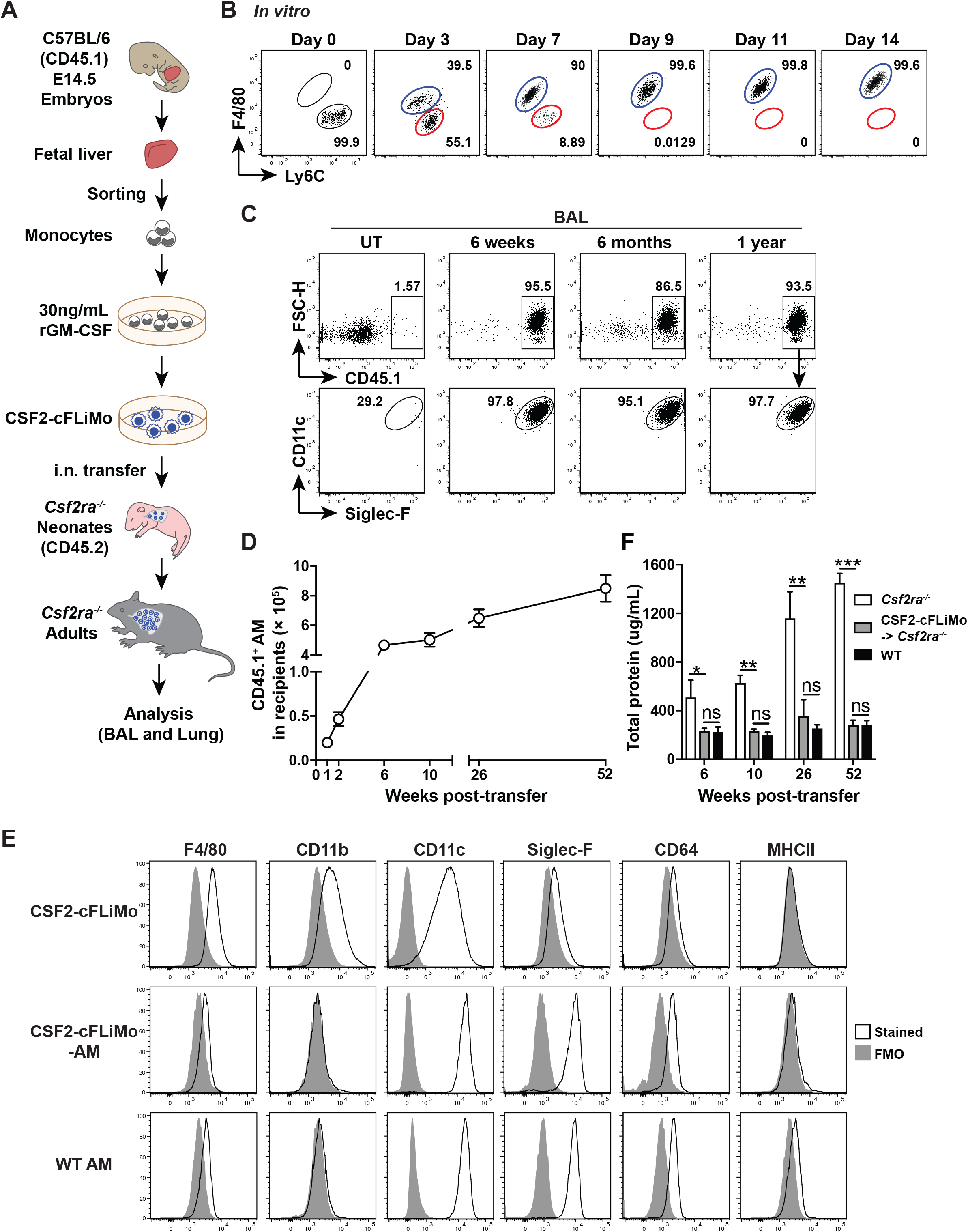
Fetal liver monocytes can proliferate *in vitro* with GM-CSF and further develop into mature functional AM *in vivo*. (A) Illustration of experimental regime. Primary monocytes from fetal liver were cultured *in vitro* with GM-CSF. (B) Representative dot plots for F4/80 vs Ly6C are shown for GM-CSF-cultured fetal liver monocytes (CSF2-cFLiMo) at the indicated time points, pre-gated on viable CD45^+^ single cells. (C-F) 5×10^4^ CSF2-cFLiMo generated from CD45.1 E14.5 embryos, after 2 weeks of culturing, were transferred intranasally (i.n.) to neonatal (day 0-3 after birth) CD45.2^+^ *Csf2ra*^*-/-*^ mice and analyzed at the indicated time points. (C) Representative dot plots of donor-derived cells in the bronchoalveolar lavage (BAL) of recipients 6 weeks, 6 months and 1 year after transfer, pre-gated as viable single cells. (D) Total numbers of donor-derived AM from lung (1 and 2 weeks) or from BAL and lung (after 6 weeks) detected in recipients at the indicated time-points. (E) Representative histograms of AM signature marker expression on CSF2-cFLiMo before transfer, CSF2-cFLiMo-derived AM detected in *Csf2ra*^*-/-*^ recipient mice 6 weeks after transfer, and endogenous AM from age-matched control mice showing FMO control (grey) and specific antibodies against the indicated markers (black line). (F) Total protein in the BAL of CSF2-cFLiMo-transferred *Csf2ra*^*-/-*^, unmanipulated *Csf2ra*^*-/-*^ and WT mice. The data are representative of three independent experiments. Values show means ± SEM of three to five mice per group in D and F. Student’s t test (unpaired) was used in F: ns, not significant; *p < 0.05, **p < 0.01, ***p < 0.001, ****p < 0.0001.

TGFβR^-/-^ mice are defective in development of AM (Yu et al, 2017). Indeed, we found that addition of TGFβ increased the proliferative capacity and the yield of CSF2-cFLiMo, but did not influence the differentiation to AM-like cells (Fig. E2A, B). Notably, culture with TGFβ in the absence of GM-CSF failed to induce proliferation and resulted in death of FLiMo (Fig. E2B). Moreover, TGFβ did not change the phenotype of CSF2-cFliMo after 2-week culture (Fig. E2C).

These results demonstrate that fetal liver monocytes cultured with GM-CSF *in vitro* give rise to a stable population of cells with AM-like phenotype and GM-CSF-dependent self-renewing capacity.

### CSF2-cFLiMo develop into mature and functional AM *in vivo*

To assess whether CSF2-cFLiMo can develop to *bona fide* AM and perform AM function *in vivo*, we transferred congenically marked CSF2-cFLiMo intranasally (i.n.) to newborn *Csf2ra*^*-/-*^ mice (Fig. 1A), which lack AM (Li et al, 2020; Schneider et al, 2017). Analysis of the BAL and lung of *Csf2ra*^*-/-*^ recipients showed efficient engraftment of donor-derived cells that resemble mature CD11c^hi^Siglec-F^hi^ AM (Fig. 1C and Fig. E3A). The numbers of CSF2-cFLiMo-derived AM rapidly increased within the first 6 weeks after transfer, before reaching a relatively stable population size (Fig. 1D), similar to the kinetics during normal postnatal AM differentiation (Guilliams et al, 2013; Schneider et al, 2014). While CSF2-cFLiMo were CD11b^hi^Siglec-F^lo^ before transfer, they down-regulated CD11b and up-regulated CD11c and Siglec-F surface expression upon transfer and expansion *in vivo*, indicating that they completed their differentiation to become cells with a phenotype that is indistinguishable from AM of age-matched WT mice (Fig. 1E). Notably, CSF2-cFLiMo-derived AM were maintained in the lung for at least 1 year after transfer (Fig. 1C, D). Moreover, measurement of protein concentration in the BAL at different time points after transfer showed that CSF2-cFLiMo-AM reconstituted *Csf2ra*^*-/-*^ mice were completely protected from PAP up to one year (Fig. 1F and Fig. E3B). These results demonstrate that CSF2-cFLiMo develop into mature AM, which appear functionally equivalent to *in situ* differentiated AM.

### CSF2-cFLiMo-derived AM can self-renew *in vivo*

AM are maintained locally through self-renewal, and they are largely independent of adult hematopoiesis at steady state (Guilliams et al, 2013; Schneider et al, 2014). Serial transplantation remains the gold standard for experimental assessment of long-term repopulating and self-renewal capacity of hematopoietic stem cells (Iscove & Nawa, 1997), but similar experiments have not been done for tissue macrophages. To determine the long-term self-renewing capacity of CSF2-cFLiMo-AM, we serially transferred *in vitro* differentiated CSF2-cFLiMo or *ex vivo* isolated AM from adult WT mice into neonatal *Csf2ra*^*-/-*^ mice. After 6 weeks, donor-derived AM were isolated and transferred into a second group of neonatal *Csf2ra*^*-/-*^ recipients (Fig. 2A). Following secondary transfer, CSF2-cFLiMo-AM and *ex vivo* AM-derived AM (AM-AM) again fully restored the AM compartment of *Csf2ra*^*-/-*^ recipients and their number and phenotype was comparable to that of AM in the BAL and lung of unmanipulated WT mice (Fig. 2B-D). These results demonstrate that CSF2-cFLiMo-AM have a self-renewal capacity *in vivo*, which is comparable to *bona fide* AM.

**Figure 2.**
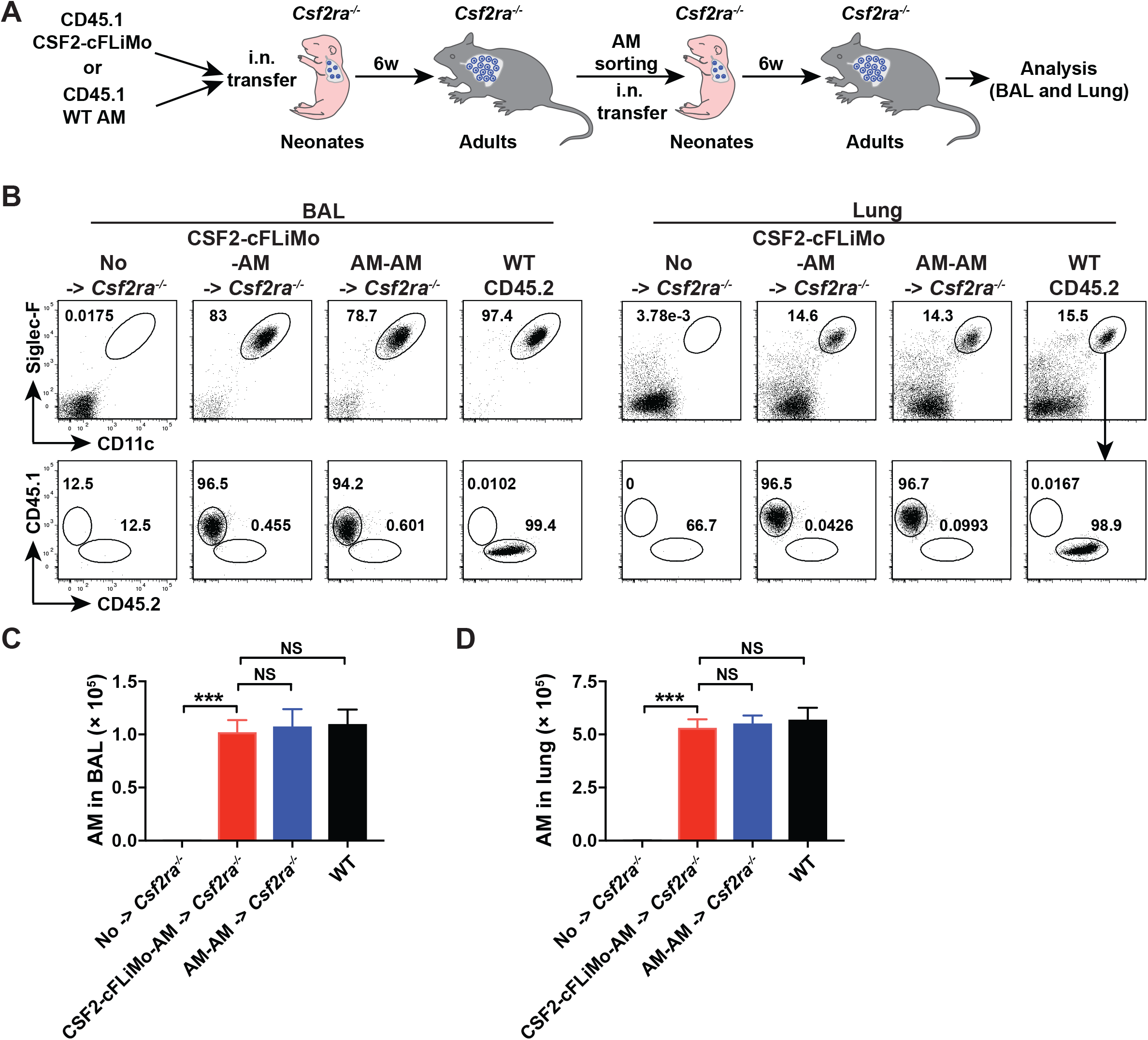
CSF2-cFLiMo-derived AM have ability to self-renew *in vivo*. (A) Illustration of experimental regime. 5×10^4^ of CD45.1 CSF2-cFLiMo generated from E14.5 embryos after 2-week culture or CD45.1 AM sorted from WT adult mice were transferred i.n. to neonatal CD45.2 *Csf2ra*^*-/-*^ mice. After 6 weeks, donor-derived AM were sorted and 5×10^4^ of cells were transferred i.n. to new neonatal *Csf2ra*^*-/-*^ mice. BAL and lung were analysed 6 weeks after second-round transfer in B-D. (B) Representative dot plots showing the phenotype of CD45.1 donor-derived AM in the BAL and lung, pre-gated as viable CD45^+^ single cells. (C-D) Numbers of donor-derived AM and WT AM in the BAL (C) and lung (D). Age-matched *Csf2ra*^*-/-*^ and CD45.2 WT mice were included as negative and positive controls, respectively. The data are representative of three experiments. Values show means ± SEM of three to four mice per group. ANOVA (one way) was used in C-D: ns, not significant; *p < 0.05, **p < 0.01, ***p < 0.001, ****p < 0.0001.

### CSF2-cFLiMo acquire AM-specific transcriptional signature *in vivo*

To reveal the gene expression programs that are associated with particular AM differentiation stages, we compared the transcriptomes of CSF2-cFLiMo, CSF2-cFLiMo-derived AM to *ex vivo* AM and E14.5 fetal liver monocytes (Fig. 3A). Principal-component analysis (PCA) and matrix clustering, based on all detected genes, revealed that CSF2-cFLiMo-derived AM displayed substantial different gene expression profiles compared to fetal liver monocytes, and CSF2-cFLiMo, while they were closely related and clustered together with *ex vivo* AM (Fig. 3B-C). CSF2-cFLiMo-AM and *ex vivo* AM expressed low levels of several well-known monocyte markers, including *Ly6c1* (Ly-6C), *Fcgr1* (CD64), *Itgam* (CD11b) (Fig. 3D). Moreover, the relative mRNA expression of several established AM markers, including *Marco, Pparg, Itgax* (CD11c), *MerTk, Cd14, Fcgr2b* (CD32), *Siglec5* (Siglec-F) and *Chil3* (Ym1) was similar between CSF2-cFLiMo-AM and *ex vivo* AM (Fig. 3D).

**Figure 3.**
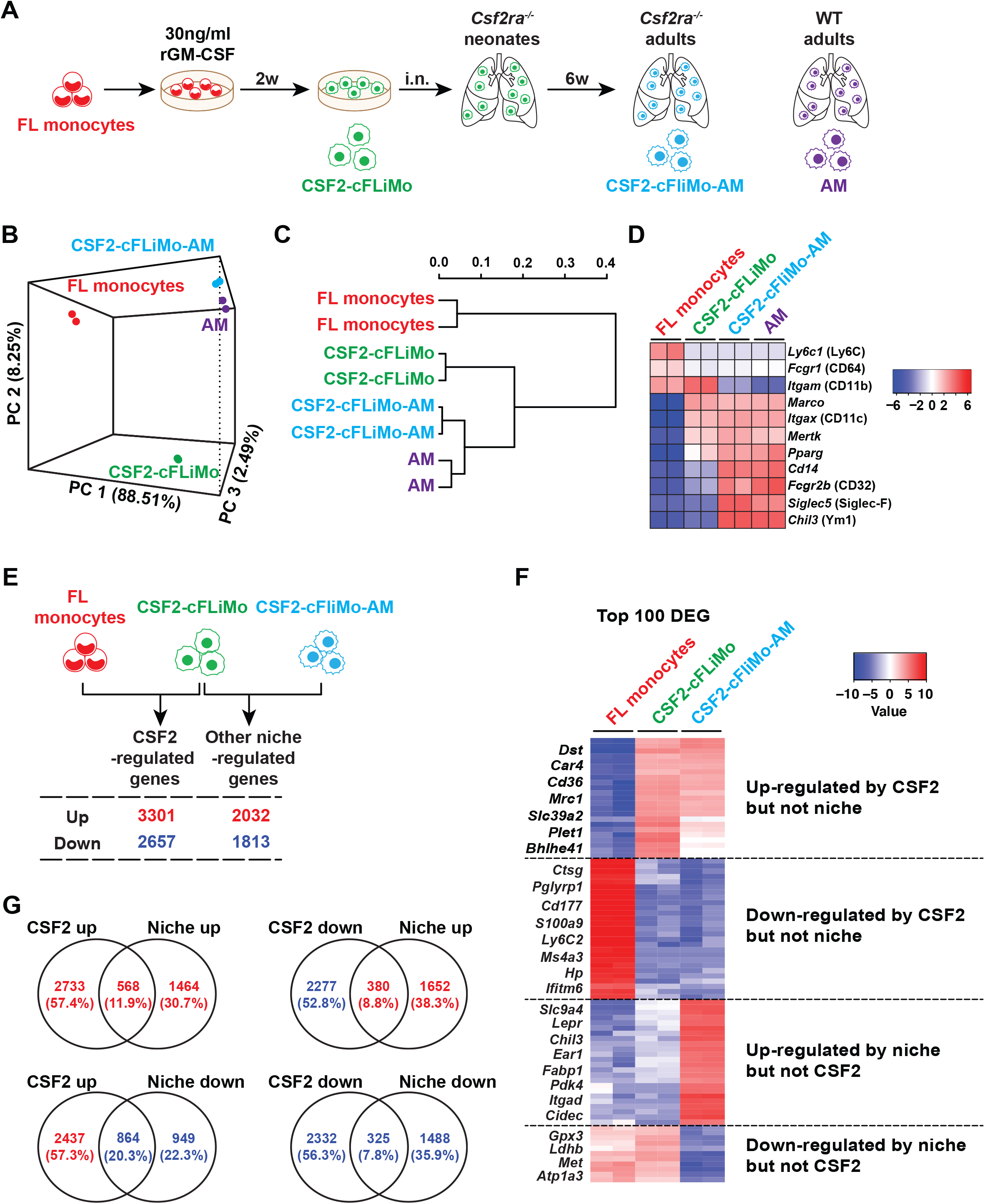
Gene expression profiles of transferred CSF2-cFLiMo in *Csf2ra*^*-/-*^ mice. (A) Illustration of experimental regime. Primary monocytes from E14.5 fetal liver were sorted using flow cytometry. CSF2-cFLiMo were collected after 2 weeks. CSF2-cFLiMo were transferred i.n. to neonatal *Csf2ra*^-/-^ mice. Mice were analysed 6 weeks after transfer and CSF2-cFLiMo-derived mature AM (CSF2-cFLiMo-AM) were sorted from the BAL as viable CD45^+^CD11c^hi^F4/80^+^Siglec-F^hi^CD11b^lo^ cells. AM from 6-week-old WT mice were sorted as control. RNA sequencing was performed (two biological replicates per group). (B) Principal component analysis (PCA) and (C) matrix clustering of the transcriptomes of all samples are shown. (D) Heat maps showing expression of monocyte and AM markers. (E) The numbers of upregulated and downregulated genes by CSF2 or niche. (F) Heat map showing the top 100 DEG and reprehensive genes of CSF2 and niche regulated. (G) Venn diagram of differentially expressed genes. Intersections of CSF2-upregulated or CSF2-downregulated vs niche-upregulated or niche-downregulated genes. The absolute gene numbers and percentages in the intersections are shown.

Next, we compared differentially expressed genes (DEG) (fold change > 2) to signature genes defined for monocytes and lung macrophages (Lavin et al, 2014). More than 60% of monocyte signature genes were downregulated, while 60% of AM signature genes were upregulated in in fetal liver monocytes cultured with GM-CSF *in vitro* (Fig. E4A), suggesting that these AM signature genes are directly or indirectly regulated by GM-CSF. Interestingly, the remaining 40% AM signature genes were up-regulated in CSF2-cFLiMo upon transfer and maturation *in vivo* (Fig. E4B).

The *in vitro* culture of precursors followed by *in vivo* transfer described here creates a model to study GM-CSF and other niche factors during AM development. Comparison of the CSF2-cFLiMo and fetal liver monocyte transcriptomes revealed 3301 upregulated and 2657 downregulated genes, which were dependent on GM-CSF (Fig. 3E). Similarly, comparing the transcriptomes of CSF2-cFLiMo-AM to CSF2-cFLiMo, we found 2032 genes that were upregulated and 1813 genes that were down-regulated by niche factors (Fig. 3E). The representative genes of the top 100 DEG regulated by CSF2 or niche factors are listed in Fig. 3F. Only a minor fraction of CSF2-upregulated genes was further upregulated (11.9%) or downregulated (20.3%) by niche factors (Fig. 3G). Similarly, 7.8% and 8.8% of CSF2-downregulated genes were further downregulated or upregulated by niche factors, respectively (Fig. 3G). These results showed that the majority of genes were separately regulated by CSF2 and additional niche factors. More than 60% of the gene expression changes were driven by CSF2 (Fig. 3G), indicating its major contribution to AM development.

Taken together, our results show that culture of fetal liver monocytes with GM-CSF *in vitro* results in AM-like precursors, which can accomplish full AM differentiation driven by alveolar niche factors upon intranasal transfer to AM-deficient neonates.

### CSF2-cFLiMo-derived AM are functional in homeostasis and during infection

Overall, our studies demonstrate that CSF2-cFLiMo-AM were functionally equivalent to naturally differentiated AM. To determine the number of donor cells required to fully reconstitute the AM compartment of *Csf2ra*^*-/-*^ mice, we titrated the number of transferred CSF2-cFLiMo (Fig. 4A). Transfer of a minimum of 5×10^4^ CSF2-cFLiMo to neonatal *Csf2ra*^*-/-*^ mice resulted in AM numbers in adult recipients that were comparable to unmanipulated WT mice (around 5×10^5^) (Fig. 4B) and protected mice from PAP (Fig. 4C). We have previously established that around 10% of primary fetal liver monocytes supplied intranasally reach the lung (Li et al, 2020). Thus, CSF2-cFLiMo have expanded around 100-fold 6 weeks after transfer to *Csf2ra*^*-/-*^ neonates. Notably, extended time of CSF2-cFLiMo *in vitro* culture (i.e. 4 months) prior transfer into recipient mice did not negatively affect their differentiation and functional capacity (Fig. 4B, C). Another critical function of tissue-resident macrophages including AM is the removal of apoptotic cells (efferocytosis) (Morioka et al, 2019). We compared efferocytosis between CSF2-cFLiMo-AM in *Csf2ra*^*-/-*^ mice and AM in WT mice by intratracheal (i.t.) installation of labelled apoptotic thymocytes. CSF2-cFLiMo-AM and AM were equally potent at phagocytosing apoptotic cells from the bronchoalveolar space (Fig. 4D).

**Figure 4.**
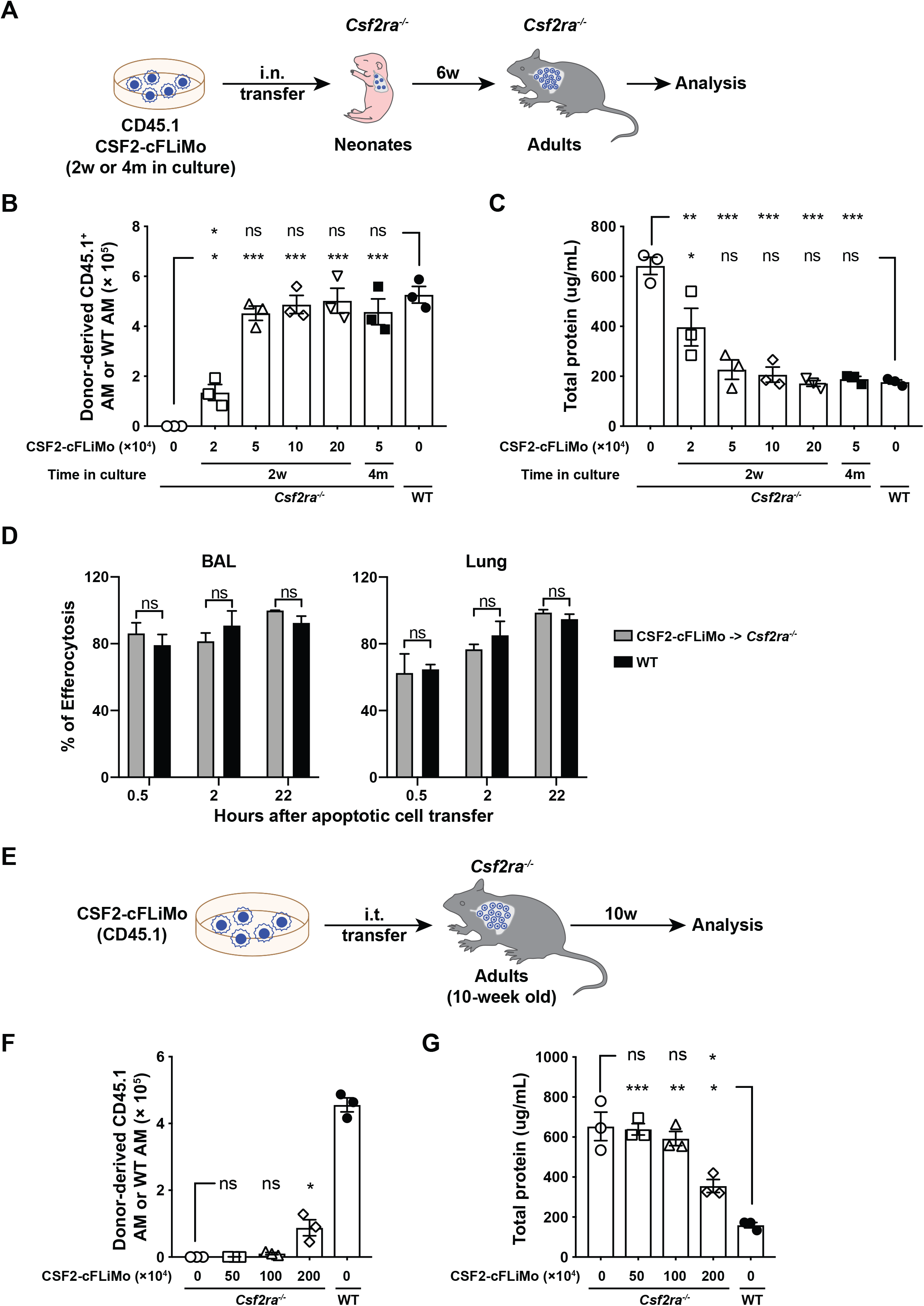
CSF2-cFLiMo-derived AM are functional in phagocytosis and efferocytosis. (A) Illustration of experimental regime. Different numbers of CD45.1 CSF2-cFLiMo after 2-week (2w) or 4-month (4m) of culturing were transferred i.n to neonatal CD45.2 *Csf2ra*^*-/-*^ mice and analysed 6 weeks later in B-C. (B) Total numbers of donor-derived AM in the BAL and lung of recipient *Csf2ra*^*-/-*^ mice or AM in the BAL and lung of WT mice. (C) Total protein levels in the BAL. (D) Efferocytosis of i.t. instilled apoptotic thymocytes by AM at the indicated time points. Values shown depict percentages of efferocytic AM. (E) Illustration of experimental regime. CD45.1 C57/BL6 CSF2-cFLiMo were generated from E14.5 embryos and cultured 2 weeks *in vitro*. Different numbers of CSF2-cFLiMo were transferred i.t. to 10-week-old adult *Csf2ra*^*-/-*^ mice and analysed 10 weeks later in F-G. (F) Total numbers of donor-derived AM in the BAL and lung of recipient *Csf2ra*^*-/-*^ mice or AM in the BAL and lung of WT mice. (G) Total protein levels in the BAL. Age-matched *Csf2ra*^*-/-*^ and WT C57BL/6 mice were included as negative and positive controls, respectively. Values show means ± SEM and the results are representative of three experiments. Student’s t test (unpaired) was used in D and ANOVA (one way) was used in B, C, F and G: ns, not significant; *p < 0.05, **p < 0.01, ***p < 0.001, ****p < 0.0001.

Next, we assessed whether CSF2-cFLiMo show therapeutic activity upon transfer into adult *Csf2ra*^*-/-*^ mice, which had already developed PAP. Adult *Csf2ra*^*-/-*^ mice were transferred i.t. with 0.5, 1 or 2 million CSF2-cFLiMo (Fig. 4E-G). Ten weeks after transfer, donor-derived AM were detectable in the BAL and lung of *Csf2ra*^*-/-*^ only in recipients transferred with 2 million cells (Fig. 4F). The protein levels in the BAL from mice transferred with 2×10^6^ cells were significantly lower when compared to naïve *Csf2ra*^*-/-*^ mice, suggesting that transferred cells were able to reduce proteinosis, although not to the level of WT mice (Fig. 4G). However, CSF2-cFLiMo-derived AM exhibited higher expression of F4/80 and CD11b, and lower expression of Siglec-F and CD64 when compared to WT AM (Fig. E5A, B), indicating that the AM phenotype was not fully recapitulated but intermediate between AM-derived from CSF2-cFLiMo transferred to neonates and AM-derived from CSF2-cFLiMo transplanted to adult mice. These results show that CSF2-cFLiMo can reproduce AM phenotype and function most adequately only when transferred to neonatal *Csf2ra*^*-/-*^ mice.

In addition to the homeostatic function, AM play an essential role in protecting influenza virus-infected mice from morbidity by maintaining lung integrity through the removal of dead cells and excess surfactant (Schneider et al, 2014). To assess the functional capacity of CSF2-cFLiMo-derived AM during pulmonary virus infection, we reconstituted *Csf2ra*^*-/-*^ neonates with CSF2-cFLiMo and infected adults 10 weeks later with influenza virus PR8 (Fig. 5A). Without transfer, *Csf2ra*^*-/-*^ mice succumbed to infection due to lung failure (Fig. 5B-E), as reported previously (Schneider et al, 2017). Notably, the presence of CSF2-cFLiMo-derived-AM protected *Csf2ra*^*-/-*^ mice from severe morbidity (Fig. 5B, C) and completely restored viability (Fig. 5D) and O_2_ saturation (Fig. 5E) compared to infected WT mice.

**Figure 5.**
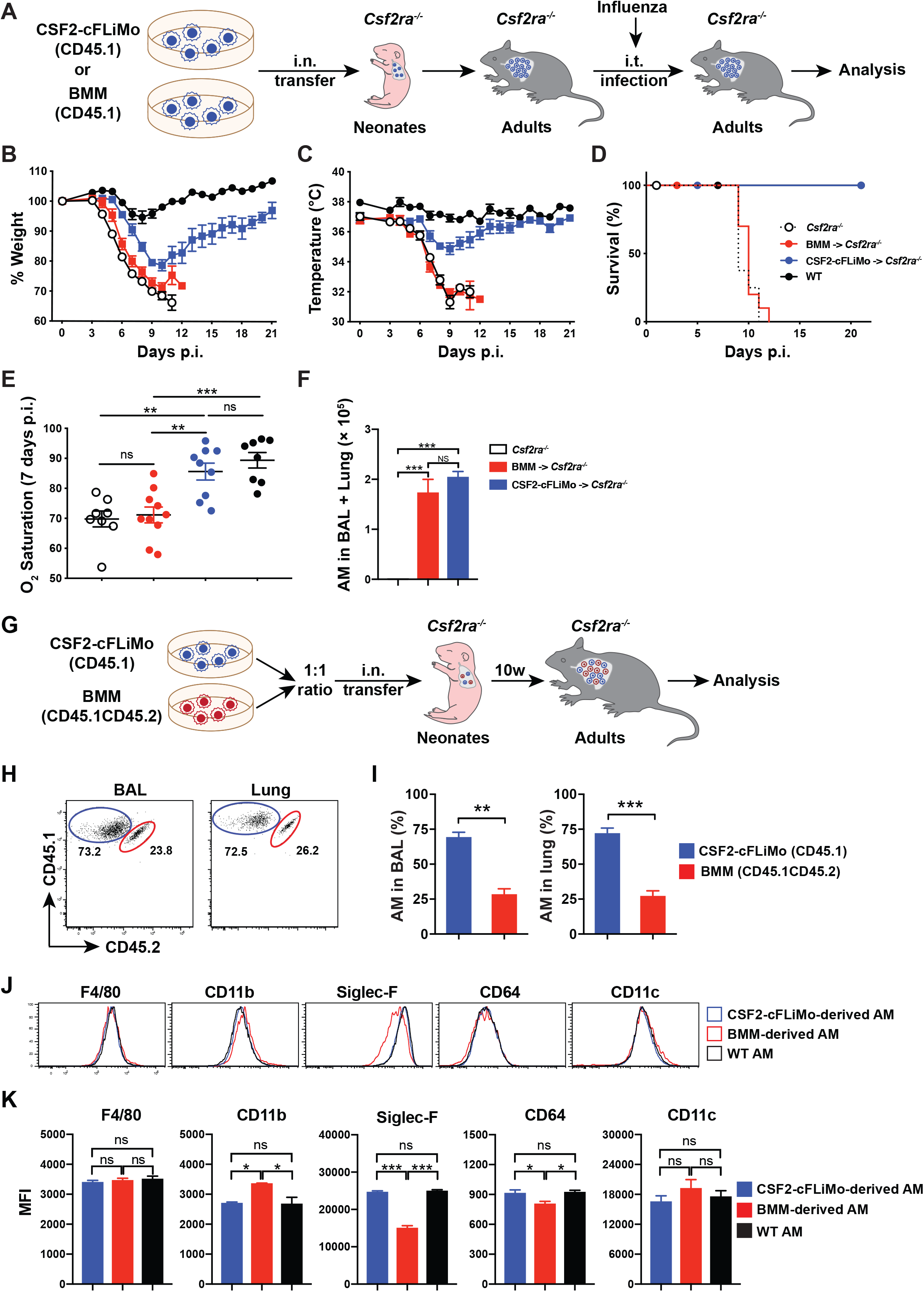
CSF2-cFLiMo, but not BMM transplantation protects *Csf2ra*^*-/-*^ mice from respiratory failure following influenza infection. (A) Illustration of experimental regime. CSF2-cFLiMo generated from E14.5 embryos after 2-week culture or BMM generated from adults after 7-day culture were i.n. transferred to neonatal CD45.2 *Csf2ra*^*-/-*^ mice. Adult recipient mice were infected i.t. with 10 pfu PR8 influenza virus and analysed in B-F. Shown are loss of body weight (B) and temperature (C), relative to the day of infection. (D) Survival curve showing the fraction of surviving animals for each day after infection. (E) Oxygen (O_2_) saturation on 7 days p.i. is shown. (F) Donor-derived AM numbers in the BAL and lung of *Csf2ra*^*-/-*^ mice 8 weeks after AM reconstitution described in Fig. 5A and prior to infection. Age-matched *Csf2ra*^*-/-*^ and WT mice were included as negative and positive controls, respectively. (G) Illustration of experimental regime. CD45.1 CSF2-cFLiMo were generated from E14.5 embryos and cultured 2 weeks *in vitro*. CD45.1CD45.2 BMM were generated from adults and cultured 7 days with M-CSF or GM-CSF *in vitro*. CSF2-cFLiMo and BMM were pooled in 1:1 ratio and transferred i.n. to neonatal CD45.2 *Csf2ra*^*-/-*^ mice and analysed 10 weeks later in H-K. (H-I) Representative dot plots (H) and percentage (I) of donor-derived AM in BAL and lung of the recipients. (J-K) Representative histograms (J) and mean fluorescence intensity (MFI) (K) of AM signature markers on CSF2-cFLiMo-and BMM-derived AM and WT AM. Values show means ± SEM and the results are representative of three experiments. Student’s t test (unpaired) was used in I and ANOVA (one way) was used in E, F and K: ns, not significant; *p < 0.05, **p < 0.01, ***p < 0.001, ****p < 0.0001.

Recent studies have shown that transplantation of BMM could decrease proteinosis in adult *Csf2rb*^*-/-*^ mice in homeostasis (Happle et al, 2014; Lachmann et al, 2014; Suzuki et al, 2014). To directly compare the capacity of BMM and CSF2-cFLiMo in AM development, we transferred M-CSF-derived BMM and CSF2-cFLiMo at a 1:1 ratio into neonatal *Csf2ra*^*-/-*^ recipient mice (Fig. 5G). Analysis AM compartment 10 weeks after transfer showed that more than 70% were derived from CSF2-cFLiMo (Fig. 4H, I) and that their phenotype (CD11b^lo^Siglec-F^hi^) closely resembled that from genuine AM, while the remaining 30% BMM-derived AM showed differences in CD11b, Siglec-F, and CD64 surface levels (Fig. 4J, K). Similar results were shown when transferring GM-CSF-derived BMM and CSF2-cFLiMo at a 1:1 ratio into neonatal *Csf2ra*^*-/-*^ recipient mice (Fig. 5G). However, transfer of BMM to *Csf2ra*^*-/-*^ neonates in the absence of competing CSF2-cFLiMo resulted in AM numbers comparable to resident AM in untreated mice (Fig. 5F). To study the function of BMM-derived AM during pulmonary virus infection, we infected *Csf2ra*^*-/-*^ recipients containing BMM-AM with influenza virus PR8 (Fig. 5A). In contrast to CSF2-cFLiMo-transferred *Csf2ra*^*-/-*^ mice, the presence of BMM-derived-AM was of no advantage and mice succumbed to infection comparable to *Csf2ra*^*-/-*^mice devoid of AM (Fig. 5B-E). These results indicate that CSF2-cFLiMo-derived AM but not BMM-derived AM protect from influenza-induced morbidity and mortality.

### Major and minor histocompatibility differences of CSF2-cFLiMo result in transplant rejection

Our results suggest that CSF2-cFLiMo could be used as an elegant *in vitro* and *in vivo* system to study the development and function of AM. To gain more information about the potential of this system, we next evaluated the MHC compatibility in an allogeneic CSF2-cFLiMo transfer. We generated CSF2-cFLiMo from BALB/c E14.5 embryos and transferred them alone, or together with C57BL/6 (B6) CSF2-cFLiMo at a 1:1 ratio into neonatal *Csf2ra*^*-/-*^ mice that were on B6 background (Fig. 6A). Ten weeks later, we were unable to detect any BALB/c CSF2-cFLiMo-AM, regardless if they had been transferred alone or together with B6 CSF2-cFLiMo, indicating that complete rejection of MHC-mismatched cells (Fig. 6B, C and Fig. E6A, B). We then also compared Y chromosome compatibility by transferring GM-CSF-differentiated monocytes isolated from male neonatal (day 0-3 after birth) liver (CSF2-cNLiMo) to newborn *Csf2ra*^*-/-*^ male or female recipients (Fig. 6D). Donor-derived AM could be detected in all transferred animals regardless of recipient’s sex (Fig. 6E, F). However, female recipients displayed significantly lower AM numbers compared to male recipients, which indicates that there was only a minor Y chromosome-dependent selection. Together, our data indicate that both major histocompatibility complex and minor histocompatibility antigens have to be considered to avoid rejection of AM grafts.

**Figure 6.**
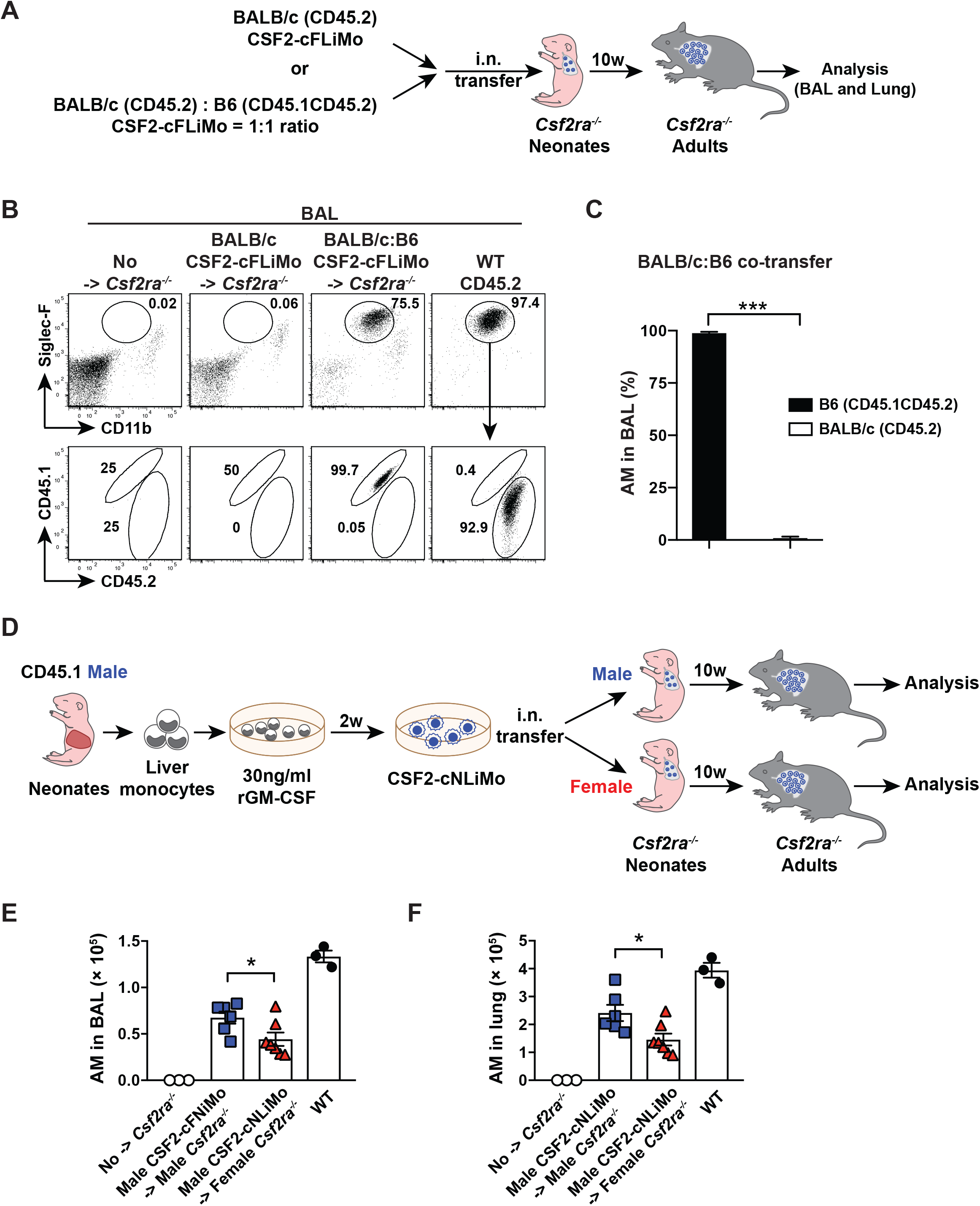
The MHC and gender compatibility in allogenic CSF2-cFLiMo transfers. (A) Illustration of experimental regime. CD45.2 BALB/c and CD45.1 B6 CSF2-cFLiMo were generated from E14.5 embryos and cultured 2 weeks *in vitro*. BALB/c CSF2-cFLiMo were transferred to neonatal CD45.2 *Csf2ra*^*-/-*^ mice (B6 background) either separately or in a 1:1 ratio with B6 CSF2-cFLiMo and analysed 10 weeks later in B-C. (B) Representative dot plots showing the phenotype of donor-derived AM in the BAL, pre-gated on viable CD45^+^ single cells. (C) Percentage of donor-derived AM in BAL of co-transferred recipients. (D) Illustration of experimental regime. CSF2-cNLiMo generated from CD45.1 WT male neonates after 2-week culture were intranasally transferred to neonatal CD45.2 *Csf2ra*^*-/-*^ mice respectively and analysed after 10 weeks in E-F. Mice were grouped according to gender. (E-F) Numbers of donor-derived AM and WT AM in the BAL (E) and lung (F) are shown. Age-matched *Csf2ra*^*-/-*^ and CD45.2 WT mice were included as negative and positive controls in B, E and F. Values show means ± SEM in C, E and F and the results are representative of three experiments. Student’s t test (unpaired) was used: ns, not significant; *p < 0.05, **p < 0.01, ***p < 0.001, ****p < 0.0001.

### Gene therapy of *Csf2ra* mutant CSF2-cFLiMo

The reconstitution of AM by transferring cultivatable precursors provides unique opportunities for genetic manipulation. To provide proof-of-concept, fetal liver monocytes were purified from E14.5 *Csf2ra*^*-/-*^ or WT embryos and transduced with a retrovirus encoding *Csf2ra-gfp* (RV^*Csf2ra-gfp*^) or control *gfp* only (RV^*gfp*^) (Fig. 7A). RV^*Csf2ra-gfp*^ transduced *Csf2ra-*deficient FLiMo expanded in the presence of GM-CSF *in vitro* (Fig. 7B) and outgrew all non-transduced cells, as indicated by the presence of almost 100% GFP^+^ cells by day 7 of culture (Fig. 7B, C). As expected, *Csf2ra-*deficient FLiMo transduced with control RV^*gfp*^ could not expand in culture (Fig. 7B, C). Notably, overexpression of *Csf2ra* in WT FLiMo provided also a slight growth advantage over non-transduced WT FLiMo *in vitro* (Fig. 7B, C). Next, we transferred RV^*Csf2ra-gfp*^ transduced *Csf2ra*-deficient cFLiMo into neonatal *Csf2ra*^*-/-*^ mice and analyzed BAL and lung 8 weeks later. Indeed, CSF2RA gene therapy of *Csf2ra-*deficient cFLiMo enabled full reconstitution of a functional AM compartment that prevented development of PAP in *Csf2ra*^*-/-*^ mice (Fig. 7D-G). Transfer of *Csf2ra*-overexpressing WT cFLiMo did not result in higher AM numbers indicating that GM-CSF bioavailability rather than receptor expression is the limiting factor. RV^*Csf2ra-gfp*^ transduced cFLiMo-derived AM were phenotypically indistinguishable from unmanipulated WT AM for multiple surface markers, including F4/80, CD11b, CD11c, Siglec-F, and MHCII (Fig 7H). These results provide a proof-of-concept that gene-modified CSF2-cFLiMo differentiate into functional *bona fide* AM, therefore allowing genetic manipulation of this important tissue macrophage compartment.

**Figure 7.**
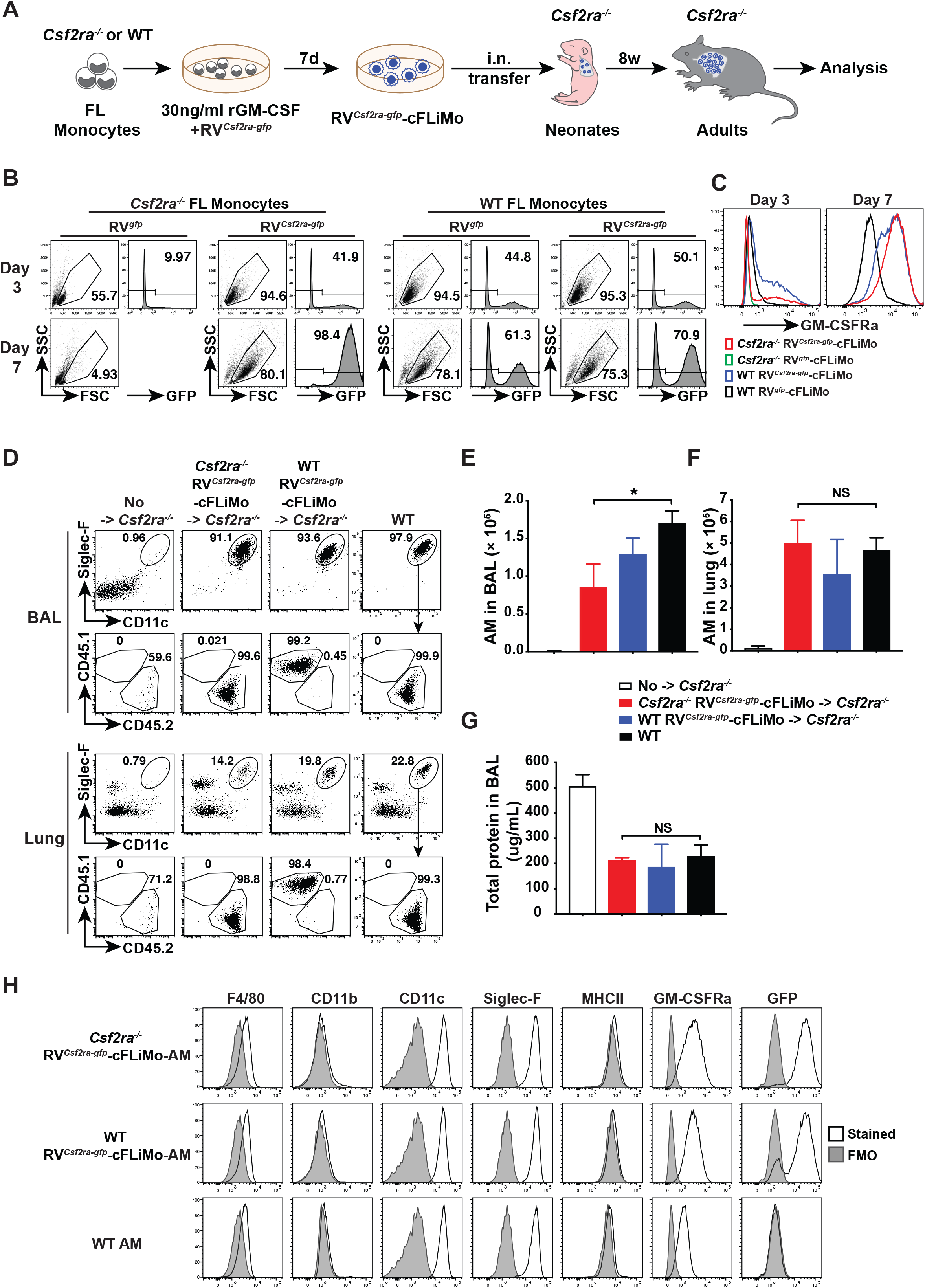
Csf2ra-reconstituted *Csf2ra*^*-/-*^ CSF2-cFLiMo can differentiate into functional AM *in vivo*. (A) Illustration of experimental regime. Fetal liver (FL) monocytes were purified from E14.5 CD45.2 *Csf2ra*^*-/-*^ or CD45.1 WT embryos and spin infected with GM-CSFRa-GFP or GFP (control) retrovirus (RV^*Csf2ra-gfp*^ or RV^*gfp*^). Cells were cultured with CSF2 for 7 days. RV^*Csf2ra-gfp*^ transduced *Csf2ra*^*-/-*^ CSF2-cFLiMo (RV^*Csf2ra-gfp*^-cFLiMo) or identically treated CD45.1 WT CSF2-cFLiMo were transferred i.n. to neonatal CD45.2^+^ *Csf2ra*^*-/-*^ mice and evaluated after 8 weeks. (B) Efficiency of spin infection (GFP^+^) and survival of cultured cells at day 3 and 7 post-infection. (C) Expression levels of GM-CSFRa on RV^*Csf2ra-gfp*^ or RV^*gfp*^ transduced *Csf2ra*^*-/-*^ or WT cFLiMo at day 3 and 7 post-infection. (D) Representative dot plots showing the phenotype of donor-derived cells in the BAL and lung, pre-gated on viable CD45^+^ single cells. (E-F) Numbers of donor-derived AM and WT AM in the BAL (E) and lung (F). (G) Total protein levels in the BAL are shown. Age-matched *Csf2ra*^*-/-*^ and CD45.2 WT mice were included as negative and positive controls in D-G, respectively. (H) Representative histograms of AM signature marker expression, GM-CSFRa and GFP on RV^*Csf2ra-gfp*^-cFLiMo-derived AM and WT AM showing FMO control (grey) and specific antibodies against indicated markers (black line). The data are representative of three experiments. Values show means ± SEM of three to four mice per group. Student’s t test (unpaired) was used in E-G: ns, not significant; *p < 0.05, **p < 0.01, ***p < 0.001, ****p < 0.0001.

## Discussion

We and others have previously shown that transfer of wild-type fetal liver or fetal lung monocytes can restore defective AM development in Csf2ra-deficient or Csf2rb-deficient mice (Guilliams et al, 2013; Li et al, 2020; Schneider et al, 2017; Schneider et al, 2014; van de Laar et al, 2016). In the present study, we describe a two-step AM differentiation model, which in a first step allows the generation of a homogenous and stable immature AM-like population (i.e. CSF2-cFLiMo) from murine fetal liver monocytes that can be maintained long-term in culture in the presence of GM-SF. In a second step, upon pulmonary transplantation, CSF2-cFLiMo expand and complete differentiation to *bona fide* AM. Indeed, low numbers of CSF2-cFLiMo completely restored the AM pool of Csf2ra-deficient mice within 6 weeks and were stably maintained for at least 1 year, indicating that they efficiently occupied empty AM niches and possess a long-term self-renewing capacity, which we have proven by serial transplantation. While serial transplantation is the gold standard for proof of long-term repopulation and self-renewal capacity of hematopoietic stem cells, according to our knowledge, to date it has not been applied to tissue macrophages.

Studies in knockout mice have shown that GM-CSF and TGF-β are essential for AM development mainly by induction of the transcription factor PPARγ and its target genes (Guilliams et al, 2013; Schneider et al, 2014; Yu et al, 2017). Our FLiMo *in vitro* cultures suggest that GM-CSF is sufficient for AM growth and differentiation of AM precursors, while TGF-β further promotes their proliferative capacity. TGF-β alone was insufficient to grow FLiMo *in vitro*. Notably, it has been suggested that fetal liver cells can be grown long-term in the presence of GM-CSF *in vitro* (Fejer et al, 2013). However, in contrast to fetal liver monocytes, fetal liver macrophages, which are derived from primitive yolk sac macrophages, could not be expanded *in vitro*, despite the presence of Csf2ra and Csf2rb chains, consistent with their poor capacity to restore AM compartment upon neonatal transfer to Csf2ra-deficient mice (Li et al, 2020).

Even though FLiMo upregulated a large panel of AM signature genes in the presence of GM-CSF *in vitro*, PCA analysis of the transcriptome showed a difference to that of mature AM. However, upon pulmonary transfer, CSF2-cFLiMo completed differentiation into *bona fide* AM indicating that lung tissue instructs terminal differentiation of AM. Thus, the two-step model allows to separate the transcriptional regulation induced by GM-CSF from other niche factors provided by the lung microenvironment during AM development. Further studies using this model will help to understand transcriptional regulation during AM development.

Notably, when CSF2-cFLiMo were transplanted to the lung of adult C*sf2ra*-deficient recipients, Siglec-F upregulation and CD11b downregulation occurred less efficiently as compared to pulmonary transplantation to neonates, indicating that CSF2-cFLiMo transplantation is more efficient in neonates than in adults. We can think of two possible reasons. Firstly, the occurrence of proteinosis in adult *Csf2ra*^*-/-*^ mice might obstruct the contact between transplanted cells and lung epithelial cells, which is important for the survival and engraftment of transplanted cells. Secondly, the neonatal lung niche could more efficiently promote AM maturation compared to the adult lung niche. Thus, the macrophage transplantation treatment for hereditary PAP patients would be more efficient during the neonatal or childhood stage before development of severe proteinosis. Importantly, we showed that the lung alveolus is not an immune privileged site implicating that transplantation AM-like precursors to MHC- and ever gender-mismatched recipients results in rejection. Thus, our data indicate that both MHC and Y chromosome compatibility require consideration when transferring CSF2-cFLiMo precursors.

M-CSF-derived BMM have been widely used to study macrophage biology *in vitro*, although they poorly recapitulate the heterogenous phenotypic and functional features of genuine resident macrophage subsets that are present in every tissue. Despite, M-CSF-derived BMM or GM-CSF-derived BMM can differentiate to AM upon pulmonary transfer to *Cs2rb-*deficient mice (Suzuki et al, 2014). Furthermore, induced pluripotent stem cell (iPSC)-derived macrophages originating from primitive macrophages in the mouse yolk sac or from human blood CD34^+^ cells have been proposed as models for pulmonary macrophage transfer therapies (Buchrieser et al, 2017; Mucci et al, 2018; Takata et al, 2017). So far, the capacity to prevent alveolar proteinosis has served as the only functional parameter for quality control in all of these approaches, except in one study that included phagocytosis of bacteria (Neehus et al, 2018).

AM develop from fetal monocytes independent of yolk sac primitive macrophages and bone marrow precursors in steady state (Hoeffel et al, 2015; Yona et al, 2013). Several studies showed recently that AM derived from primitive yolk-sac derived macrophages or from bone marrow monocytes are different in metabolism and function (Aegerter et al, 2020; Gibbings et al, 2015; Machiels et al, 2017; Misharin et al, 2017). In fact, comparing AM derived from either CSF2-cFLiMo or BMM, we found that the latter possess impaired capacity to engraft, expand, and acquire a *bona fide* AM phenotype and function including protection of *Csf2ra*^*-/-*^ mice from mortality and respiratory failure during influenza infection. These results demonstrate that catabolism of accumulated proteins in the BAL is insufficient to assess the efficacy of pulmonary macrophage transplantation therapies and that GM-CSF cultured FLiMo are superior to AM-like cells derived from bone marrow or blood cultures. CSF2-cFLiMo-derived AM prevented PAP, performed efferocytosis of apoptotic cells, and protected from fatal respiratory viral infection, indicating that they acquired the broad functional spectrum of *bona fide* AM.

CSF2-cFLiMo generated from wild-type or gene-deficient mice could be used as a high-throughput screening system to study AM development *in vitro* and *in vivo*. Our model is suitable to study the relationship between AM and lung tissue, as well as the roles of specific genes or factors in AM development and function. Furthermore, CSF2-cFLiMo can overcome the limitation in macrophage precursor numbers and be used as a therapeutic approach for PAP disease or in other macrophage-based cell therapies including lung emphysema, lung fibrosis, lung infectious disease and lung cancer (Byrne et al, 2016; Lee et al, 2016; Wilson et al, 2010). Finally, genetically modified and transferred CSF2-cFLiMo might facilitate the controlled expression of specific therapeutic proteins in the lung for disease treatment, and therefore, could represent an attractive alternative to non-specific gene delivery by viral vectors.

## Materials and Methods

### Mice

C57BL/6 CD45.2, congenic CD45.1 and, BALB/c mice were originally from the Jackson Laboratory (Bar Harbor, Maine, USA). *Csf2ra*^*-/-*^ mice were established in our laboratory (Schneider et al, 2017). All mice were housed and bred under specific pathogen-free conditions in individually ventilated cages in a controlled day-night cycle at the ETH Phenomics Facility. Mice used for experiments were 6-10 weeks (adults) of age, unless otherwise stated. All animal experiments were performed according to the guidelines (Swiss Animal Protection Ordinance (TschV) Zurich) and Swiss animal protection law (TschG) and had been approved by the local animal ethics committee (Kantonales Veterinaersamt Zurich).

### Timed pregnancy

Female C57BL/6 CD45.1, CD45.2, or BALB/c mice were housed together with matching male mice overnight. The vaginal plug was checked on the next day and was designated as embryonic day 0.5 (E0.5).

### Pulmonary cell transplantation

Neonatal (day 0-3 after birth) *Csf2ra*^*-/-*^ recipient mice were transferred intranasally (i.n.) with different numbers of cells in 10 μL endotoxin-free PBS. For competitive transfer experiments, 25,000 cells from each origin were mixed and transferred. Adult *Csf2ra*^*-/-*^ recipient mice were transferred intratracheally (i.t.) with different numbers of cells in 50 μL endotoxin-free PBS.

### Cell suspension preparation

Mice were euthanized by overdose (400 mg/kg body weight) of sodium pentobarbital by i.p. injection. The lungs were washed three times with 0.4 ml of ice-cold PBS containing 2 mM EDTA through an intratracheal cannula only when mice were older than 4 weeks. BAL fluid was collected and cells were harvested by centrifugation. Lungs were removed after perfusion with ice-cold PBS. Pregnant females were sacrificed by CO_2_ asphyxiation. Fetal livers were removed at the indicated time points. Organs were minced and then digested at 37 °C in IMDM medium containing 2.0 mg/ml of type IV collagenase (Worthington), 0.125 mg/ml DNase I (Sigma) and 3% FCS for 45 min (lungs) or 15 min (fetal and neonatal livers) respectively, and subsequently passed through a 70-μm-cell strainer (Becton Dickinson). Ammonium-chloride-potassium (ACK) lysing buffer was used for erythrocyte lysis for all samples.

### Flow cytometry and cell sorting

Multiparameter assessment and cell sorting were performed using LSR Fortessa, BD FACS ARIA II and ARIA III (BD Biosciences) and data were analyzed with FlowJo software (TreeStar). After blocking the FcgIII/II receptors by incubation with homemade anti-CD16/32 (2.4G2), single-cell suspensions were incubated with the indicated fluorochrome-conjugated or biotinylated monoclonal antibodies in FACS buffer (PBS containing 2% FCS and 2 mM EDTA) and then washed twice before detection. Monoclonal antibodies specific to mouse CD45 (30-F11), CD11c (N418), F4/80 (BM8), CD11b (M1/70), Siglec-F (E50-2440, BD Biosciences), CD45.1 (A20), CD45.2 (104), Ly6C (HK1.4), GM-CSFRa (698423, R&D), CD64 (X54-5/7.1) and MHC class II (M5/114.15.2, eBioscience) were purchased from BioLegend unless otherwise stated. Dead cells were excluded using the live/dead marker eFluor780 (eBioscience).

### Generation of CSF2-cFLiMo

Fetal livers were harvested from CD45.1^+^ C57BL/6 embryos. Liver single-cell suspensions were prepared. Monocytes were sorted using flow cytometry. Sorted monocytes were cultured in complete RPMI, supplemented with GM-CSF (PeproTech) (30 ng/mL) *in vitro*. Cells were plated with 1×10^5^ cells/ml and sub-cultured by splitting them 1:5 every 3 days. CSF2-cFLiMo used for transplantation were cultured for 2 weeks in culture media prior to transfers unless stated otherwise.

### Generation of BMM

For the preparation of mouse BM-derived macrophages (BMM), tibias and femurs from the hind legs of adult donor mice were flushed with PBS. Bone marrow was rinsed through a 70-um cell strainer (Becton Dickinson) followed by red blood cell depletion with ammonium-chloride-potassium (ACK) lysing buffer. BMM were differentiated *in vitro* in complete RPMI, supplemented with 10 ng/ml M-CSF (PeproTech) or 30ng/ml GM-CSF (PeproTech). Medium was replaced on day 3 and day 6. Adherent cells were harvested and used as mature BMM on day 7 or 8.

### Assessment of total protein

Total protein concentrations in BAL fluid were detected by Pierce BCA Protein Assay Kit according to the manufacturer’s instructions (Thermo Scientific).

### RNA sequencing

100,000 cells of indicated populations were collected into TRIzol (Life Technologies). Phase separation was achieved with the addition of chloroform (Sigma), and total RNA was precipitated from the aqueous layer with isopropanol (Sigma) using glycogen (Roche) as a carrier. RNA samples were sent to the Functional Genomics Center Zurich, where the RNA sequencing was performed. The TruSeq RNA Stranded sample kit (Illumina) was used to construct the sequencing libraries. In brief, total RNA samples (100 ng) were poly (A) enriched and reverse-transcribed into double-stranded cDNA, and TruSeq adapters were then ligated to double-stranded cDNA, then fragments containing TruSeq adapters on both ends were selectively enriched with PCR and subsequently sequenced on the Illumina Nova Seq. The fragments were mapped to the ensemble mouse reference genome GRCm38 (Version25.06.2015) using the STAR aligner (Dobin et al, 2013). For normalization, the read counts were scaled with the use of the trimmed mean of M-values (TMM) method proposed by Robinson and Oshlack (Robinson & Oshlack, 2010). Principal-component analysis, matrix clustering and heatmap were generated using R.

### Efferocytosis of apoptotic cells

Thymocytes were isolated from mice and apoptosis was induced by exposure to 60 mJ/cm^2^ UV radiation (Spectrolinker XL-1500; Spectronics Corporation). After 2 h incubation at 37 °C in IMDM + 10% FCS, cells were labeled with 5 mM eFluor670 (eBioscience) according to the manufacturer’s instructions, washed extensively with IMDM + 10% FCS and PBS. Then apoptotic cells (5 million cells in 50 μL PBS) were delivered i.t. to recipient mice. 0.5, 2 and 22 h after administration, efferocytosis by AM in the BAL and lung was assessed by flow cytometry.

### Influenza viral infection

Influenza virus strain PR8 (A/Puerto Rico/34, H1N1) was originally provided by J. Pavlovic, University Zurich. For infections, the mice were anesthetized and intratracheally (i.t.) inoculated with indicated doses of virus in 50 ul endotoxin-free PBS. Temperature and weight of mice were monitored daily and animals were euthanized if they fulfilled the severity criteria set out by institutional and cantonal guidelines.

### Measurement of oxygen saturation

The MouseOx™ Puls-oximeter (Starr Life Sciences) was used to measure oxygen (O_2_) saturation in influenza-infected mice on day 7 post infection. The depilatory agent was applied to the neck of mice 2-3 days prior to measurement to remove hair. Mice were sedated with 2.5mg/kg intraperitoneal Midazolam (Roche) 0.5-1 hour before measurement. The sensor clip was placed on the neck and O_2_ saturation was measured each second over 3-5 min per mouse. Data shown is the average value of each mouse.

### RV-reconstitution of CSF2RA gene

Two retroviral constructs based on moloney murine leukemia virus, containing Csf2ra cDNA and GFP (RV^*Csf2ra-gfp*^) or GFP only (RV^*gfp*^), were used for transfection of retrovirus packaging cell line. Fresh viral supernatants containing non-replicating retroviruses were used for transduction of fetal liver monocytes isolated as described above from *Csf2ra*^*-/-*^ or CD45.1^+^ WT embryos. Cells were cultured in complete RPMI, supplemented with GM-CSF (30 ng/mL) for 7 days. All non-transduced fetal liver monocytes derived from *Csf2ra*^*-/-*^ embryos could not proliferate. After a week, a homogenous population of receptor-expressing cells was detected and used for transfers. For WT cells, all live cells treated with the Csf2ra-overexpressing virus were used (irrespectively of the transduction level).

### Statistical analysis

Mean values and SEM were calculated with Prism (GraphPad Software, Inc). Student’s t-test (unpaired) was used for comparing two groups, and ANOVA (one way) was used for comparing multiple groups: ns, not significant; *p < 0.05, **p < 0.01, ***p < 0.001, ****p < 0.0001.

## Acknowledgements

We thank the teams of the ETH Flow Cytometry Core Facility for cell sorting and the EPIC mouse facility for animal husbandry, Peter Nielsen for discussion and editing of the manuscript. We are grateful for research grants from SNF (310030_163443 and 310030B_182829).

## Author contributions

F.L., K.O., and M.K. designed the experiments; F.L. and K.O. performed and analyzed the experiments. L.P. analyzed RNA sequencing data. C.S. and M.K. discussed data and provided conceptualization. F.L., K.O., and M.K. wrote the manuscript.

## Notes

### Competing Interest Statement

The authors have declared no competing interest.

### Summary of Updates

There are minor revisions on Title, Abstract, Introduction, Results, Discussion Methods, Figure legends, and references.

