## Supplemental Figure Legends for "Long-term culture of fetal monocyte precursors *in vitro* allowing the generation of *bona fide* alveolar macrophages *in vivo*"

**Supplementary Figure Legends**

**Figure E1. Fetal liver monocytes differentiate into long-lived, homogeneous, AM-like cells *in vitro* in the presence of GM-CSF, related to Figure 1**

(A) Sorting strategy for primitive macrophages (pMΦ) and fetal liver monocytes from fetal liver of CD45.1 C57BL/6 embryos. Doublets and debris were excluded using FSC and SSC. fetal liver monocytes were identified as viable CD45^+^F4/80^lo^CD11b^int^Ly6C^+^ cells and pMΦ were identified as viable CD45^+^F4/80^hi^ CD11b^lo^ cells. (B) Growth curves of fetal liver monocytes and pMΦ cultured *in vitro* with GM-CSF (30 ng/mL). (C-E) Flow cytometry was used to characterize cultured cells. (C) Histograms for SSC-A, F4/80, CD11c and Siglec-F expression on F4/80^+^Ly6C^-^ macrophages at the indicated time points. (D-E) Representative dot plots of CSF2-cFLiMo from E14.5 embryos after 8, 10 and 18 weeks (D) or from E16.5, E18.5 and E20.5 embryos after 8 weeks *in vitro* (E), pre-gated on viable CD45^+^ single cells. (F) Illustration of experimental regime. CSF2-cFLiMo cultured for more than 2 weeks in the presence of GM-CSF. The effects of GM-CSF deprivation were analyzed in G-H. (G) Numbers of CSF2-cFLiMo after removing GM-CSF. (H) Phenotype of CSF2-cFLiMo 6 days after removing GM-CSF. The data are representative of three experiments. Data are shown of three mice per group in D and E and values show means ± SEM two wells per group in B and G. Student’s t test (unpaired) was used in G: ns, not significant; *p < 0.05, **p < 0.01, ***p < 0.001, ****p < 0.0001.

**Figure E2. Fetal liver monocytes culture with GM-CSF and TGFβ, related to Figure 1**

(A) Illustration of experimental regime. Monocytes from fetal liver were cultured *in vitro* with GM-CSF (30ng/mL) and/or TGFβ (0.2ng/mL). (B) Growth curves of cFLiMo. (C) Phenotype of cFLiMo 14 days after culturing. The data are representative of three experiments. Values show means ± SEM two wells per group in B. Student’s t test (unpaired) was used in B to compare GM-CSF group and GM-CSF + TGFβ group: ns, not significant; *p < 0.05, **p < 0.01, ***p < 0.001, ****p < 0.0001.

**Figure E3. CSF2-cFLiMo have the potential to further differentiate into mature functional AM, related to Figure 1**

CSF2-cFLiMo generated from CD45.1 E14.5 embryos were transferred i.n. to CD45.2 *Csf2ra^-/-^* mice and analysed at the indicated time points as described in Fig. 1A. (A) Representative dot plots of donor-derived cells in the lung of recipients 6 weeks, 6 months and 1 year after transfer, pre-gated as viable single cells. (B) Appearance of BAL fluid 1 year after transfer. The data are representative of three independent experiments.

**Figure E4. Gene expression profiles of CSF2-cFLiMo *in vitro* and *in vivo*, related to Figure 3**

RNA sequencing was performed as described in Figure 3A. Bar graphs showing up-regulated and down-regulated (blue) genes from a comparison of CSF2-cFLiMo vs fetal liver monocytes (A) and CSF2-cFLiMo-derived AM vs CSF2-cFLiMo (B) (fold change > 2, P < 0.01, Fdr < 0.01) with gene signatures of monocytes and Lung MΦ derived from Lavin *et al*. (2014).

**Figure E5. Transplantation of CSF2-cFLiMo in adult *Csf2ra^-/-^* mice, related to Figure 4**

CSF2-cFLiMo were transferred i.t. to adult *Csf2ra^-/-^* mice and analysed 10 weeks later as described in Fig. 4E. (A) Representative histograms and (B) MFI of AM signature markers on CSF2-cFLiMo- derived AM and WT AM. Values show means ± SEM in B and the results are representative of three experiments. Student's t-test (unpaired) was used: ns, not significant; *p < 0.05, **p < 0.01, ***p < 0.001, ****p < 0.0001.

**Figure E6. The MHC compatibility of allogenic CSF2-cFLiMo transfer, related to Figure 6**

BALB/c CSF2-cFLiMo were transferred to neonatal CD45.2 *Csf2ra^-/-^* mice either separately or in 1:1 ratio with B6 CSF2-cFLiMo and analysed 10 weeks later as described in Fig. 6A. (A) Representative dot plots showing the phenotype of donor-derived AM in the lung, pre-gated on viable CD45^+^ single cells. (B) Percentage of donor-derived AM in lungs of co-transferred recipients. Age-matched *Csf2ra^-/-^* and CD45.2 WT mice were included as negative and positive controls in A. Values show means ± SEM and the results are representative of three experiments. Student’s t test (unpaired) was used: ns, not significant; *p < 0.05, **p < 0.01, ***p < 0.001, ****p < 0.0001.
