## Supplementary figures and images for "Long-term culture of fetal monocyte precursors *in vitro* allowing the generation of *bona fide* alveolar macrophages *in vivo*"

### Supplemental Figures

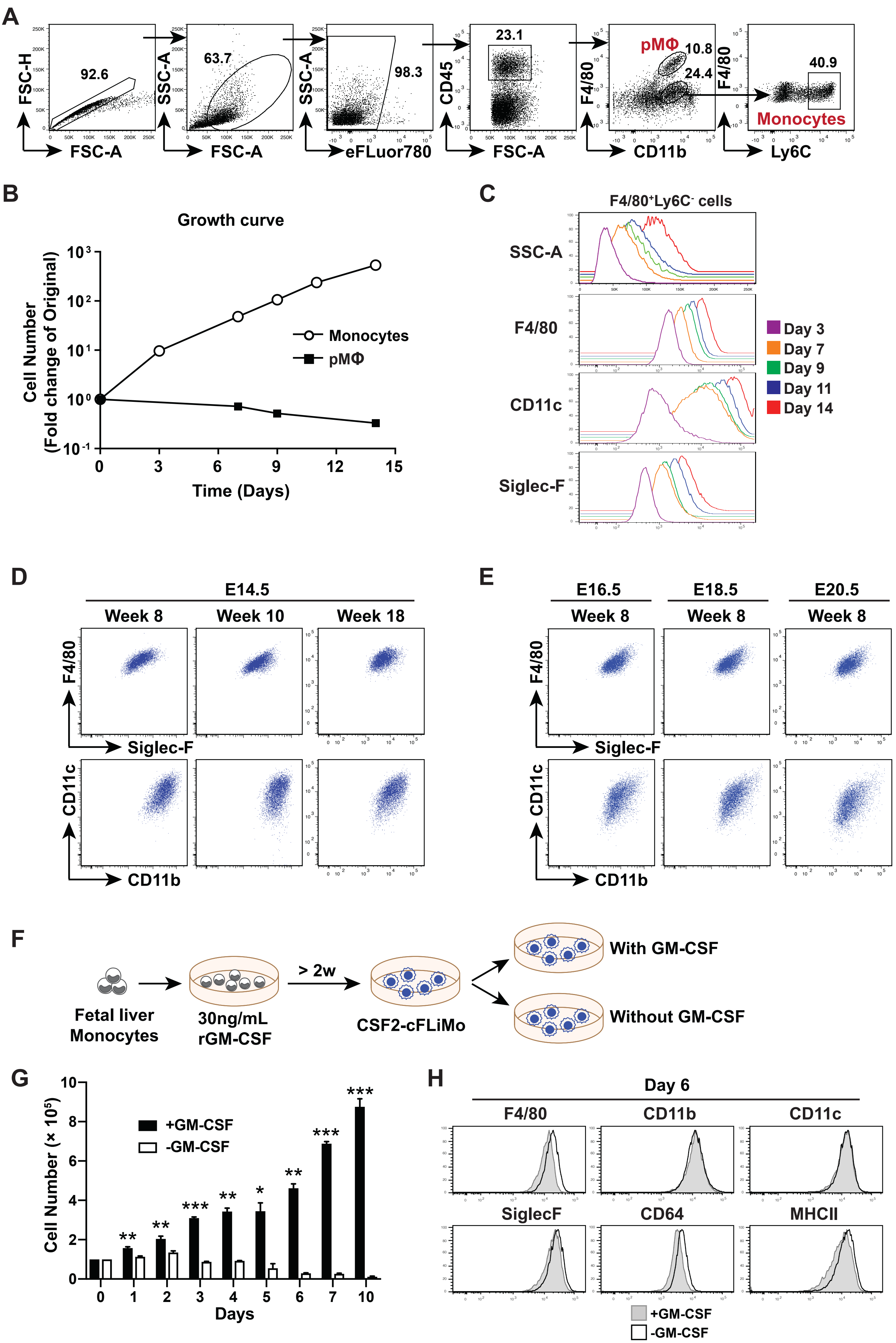

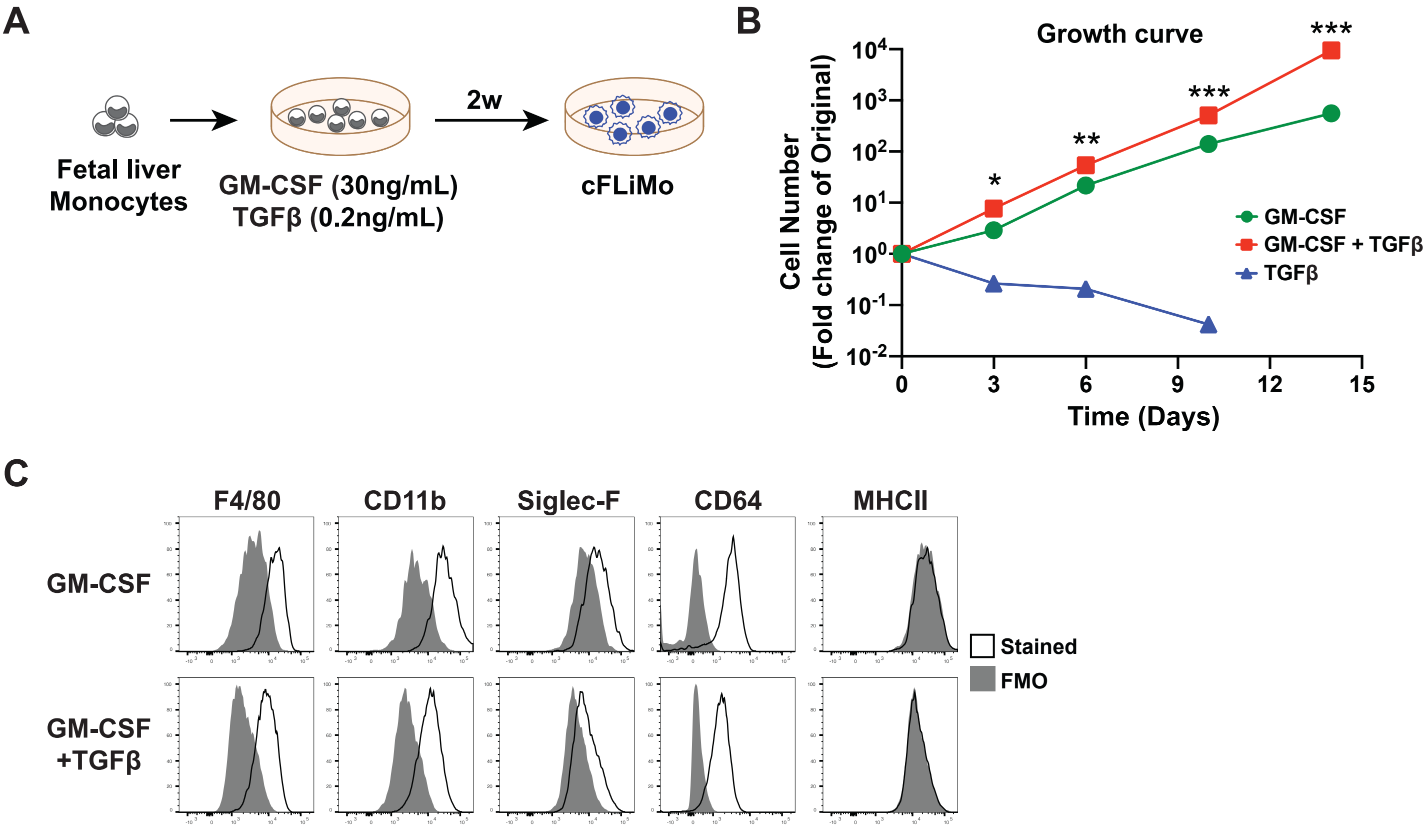

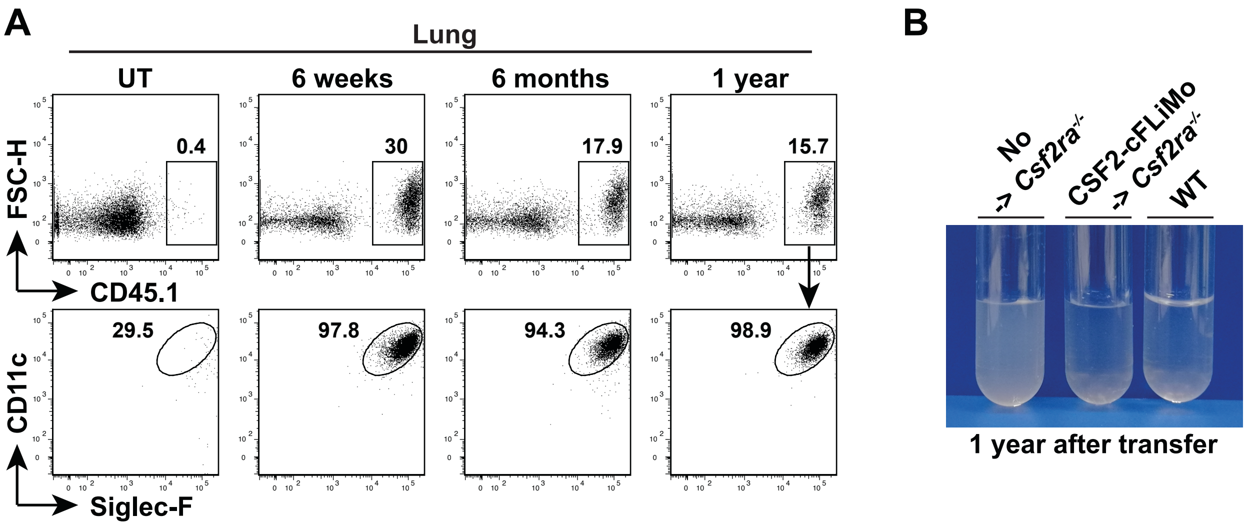

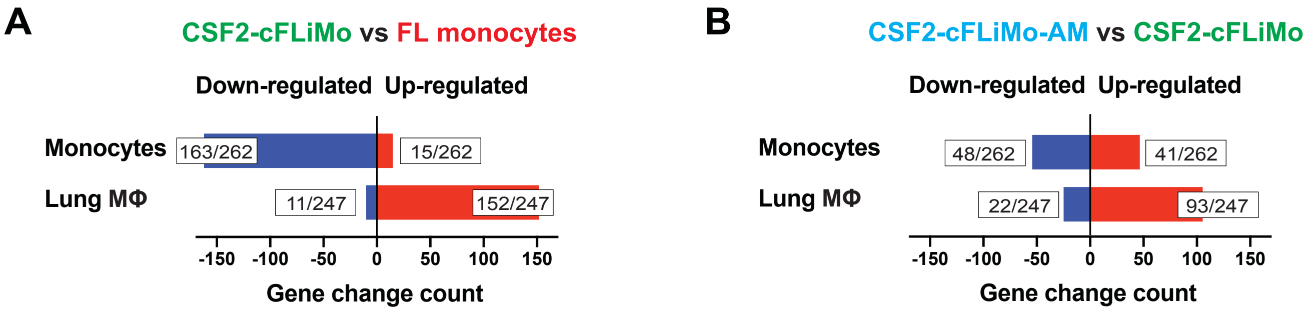

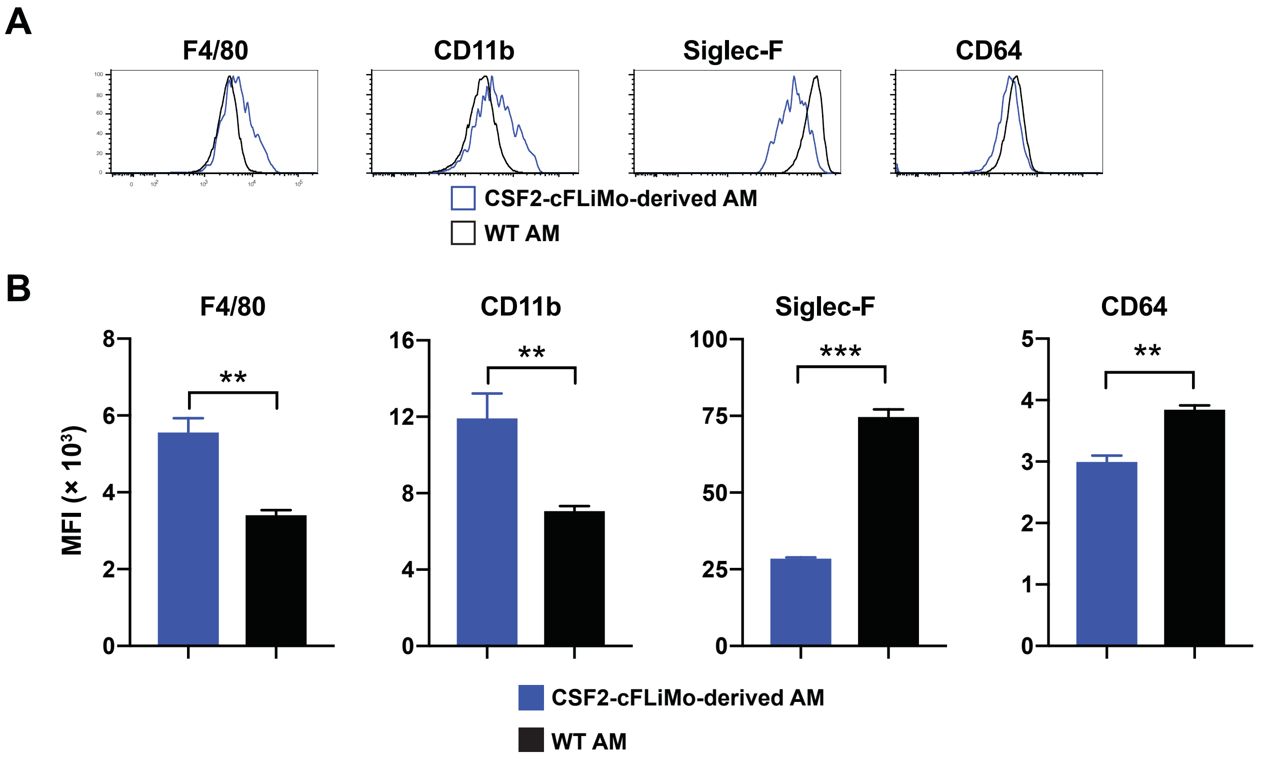

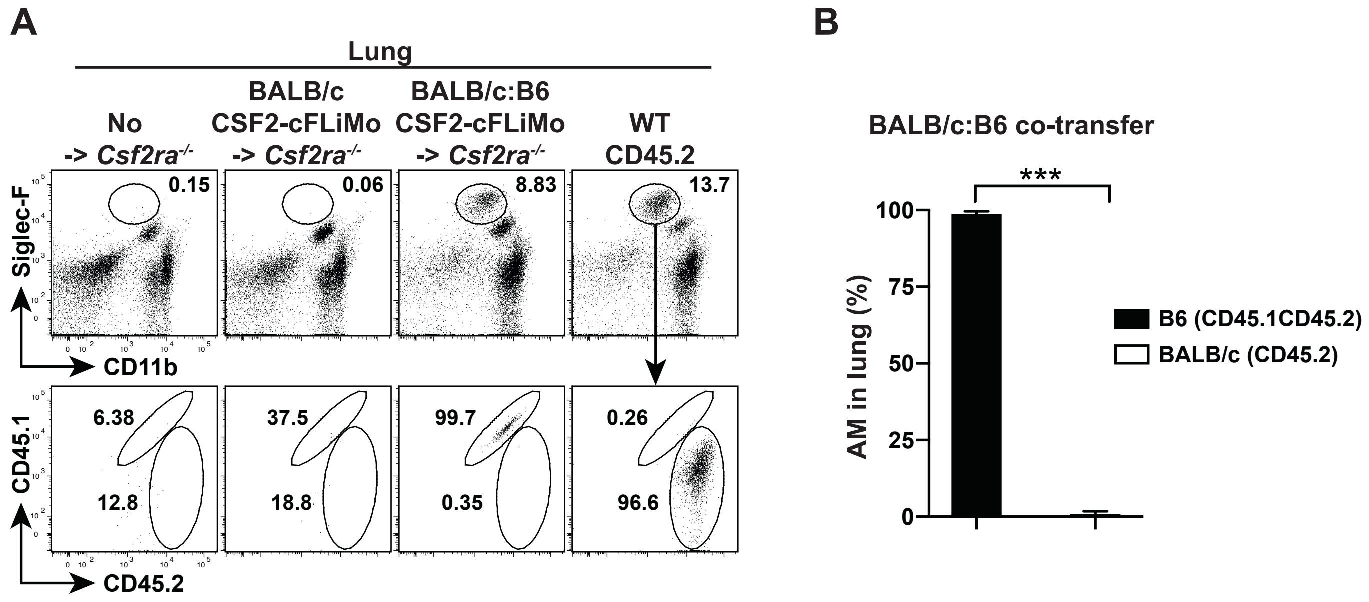
